# Examination of signatures of recent positive selection on genes involved in human sialic acid biology

**DOI:** 10.1101/137034

**Authors:** Jiyun M. Moon, David M. Aronoff, John A. Capra, Patrick Abbot, Antonis Rokas

## Abstract

Sialic acids are nine carbon sugars ubiquitously found on the surfaces of vertebrate cells and are involved in various immune response-related processes. In humans, at least 58 genes spanning diverse functions, from biosynthesis and activation to recycling and degradation, are involved in sialic acid biology. Because of their role in immunity, sialic acid biology genes have been hypothesized to exhibit elevated rates of evolutionary change. Consistent with this hypothesis, several genes involved in sialic acid biology have experienced higher rates of non-synonymous substitutions in the human lineage than their counterparts in other great apes, perhaps in response to ancient pathogens that infected hominins millions of years ago (paleopathogens). To test whether sialic acid biology genes have also experienced more recent positive selection during the evolution of the modern human lineage, reflecting adaptation to contemporary cosmopolitan or geographically-restricted pathogens, we examined whether their protein-coding regions showed evidence of recent hard and soft selective sweeps. This examination involved the calculation of four measures that quantify changes in allele frequency spectra, extent of population differentiation, and haplotype homozygosity caused by recent hard and soft selective sweeps for 55 sialic acid biology genes using publicly available whole genome sequencing data from 1,668 humans from three ethnic groups. To disentangle evidence for selection from confounding demographic effects, we compared the observed patterns in sialic acid biology genes to simulated sequences of the same length under a model of neutral evolution that takes into account human demographic history. We found that the patterns of genetic variation of most sialic acid biology genes did not significantly deviate from neutral expectations and were not significantly different among genes belonging to different functional categories. Those few sialic acid biology genes that significantly deviated from neutrality either experienced soft sweeps or population-specific hard sweeps. Interestingly, while most hard sweeps occurred on genes involved in sialic acid recognition, most soft sweeps involved genes associated with recycling, degradation and activation, transport, and transfer functions. We propose that the lack of signatures of recent positive selection for the majority of the sialic acid biology genes is consistent with the view that these genes regulate immune responses against ancient rather than contemporary cosmopolitan or geographically restricted pathogens.

## Introduction

Sialic acids are nine carbon sugars that are commonly found on the ends of glycoconjugates in deuterostome animals (Schauer 1982; Varki 2007). These molecules are involved in several biological processes, such as intercellular adhesion and signaling (Varki & Varki 2007; Kelm & Schauer 1997), and play important roles in the modulation of various aspects of the host immune system (Varki & Gagneux 2012; Pilatte et al. 1993) including activation of immune responses, leukocyte trafficking, complement pathway activation, and microbial attachment. Modification of sialic acid molecules during their biosynthesis and subsequent attachments to underlying sugars via different linkages give rise to a diverse repertoire of sialic acids (Angata & Varki 2002; Varki & Varki 2007; Cohen & Varki 2010; Hayakawa & Varki 2011).

More than 50 genes are known to be involved in various aspects of sialic acid biology in humans, and they fall into five broad functional categories (Altheide et al. 2006): 1) biosynthesis, 2) activation, transport, and transfer, 3) modification, 4) recognition, and 5) recycling and degradation (Figure 1). Biosynthesis genes are involved in assembling sialic acids from precursor molecules in the cytosol, while genes involved in the second category activate the sialic acid by attaching cytidine monophosphate to them, transport the activated sialic acids to the Golgi for attachment to glycoconjugates via multiple forms of α linkages, and transfer them to the cell surface. Activated sialic acids that are linked to the underlying sugar can be further modified by incorporation of additional molecules before the sialylated glycoconjugate is transferred to the cell surface. At the cell surface, sialic acids are recognized by specialized receptors and finally, the used sialic acids end up in the lysosome for recycling and degradation (Altheide et al. 2006).

**Figure 1.**
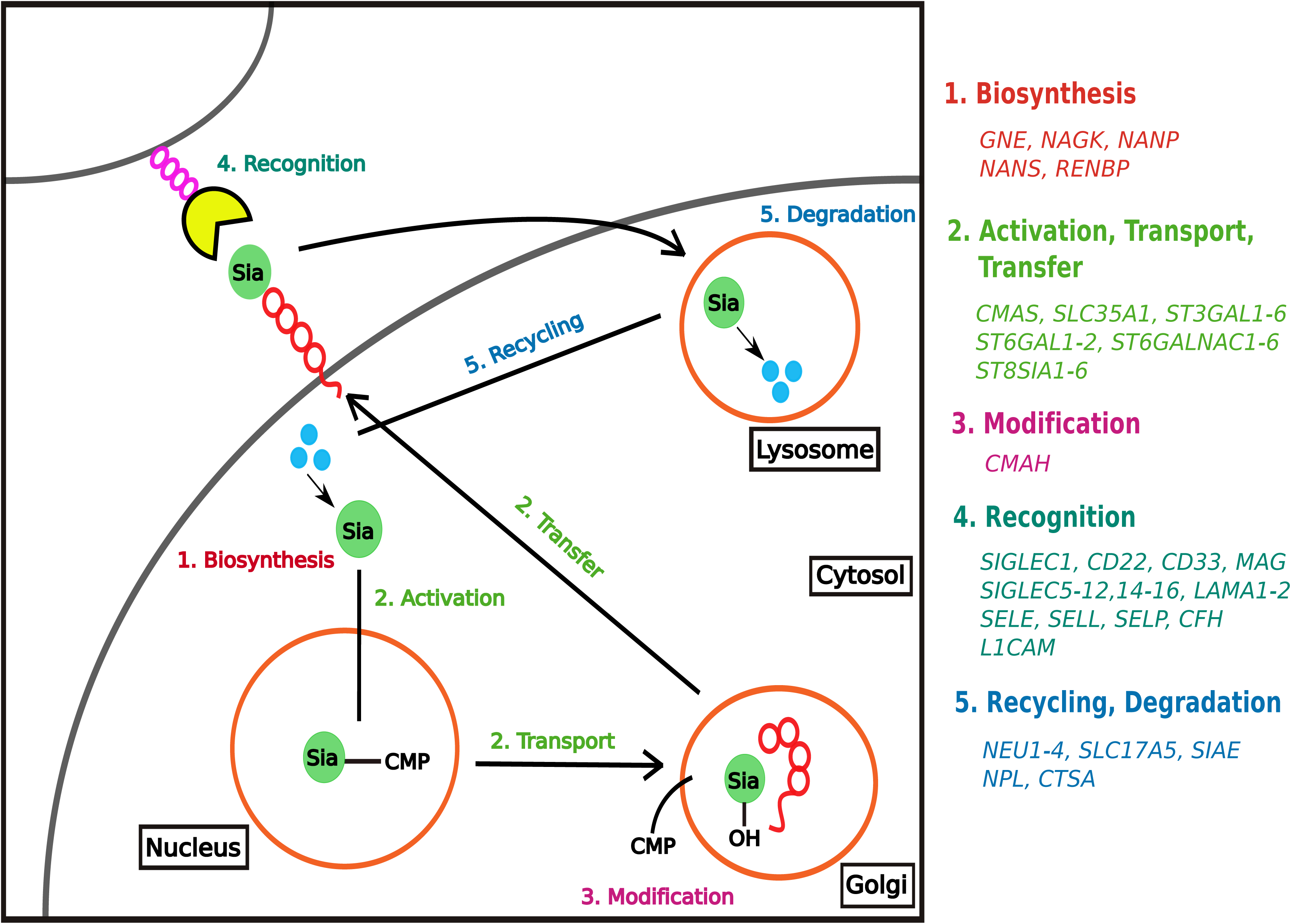
Summary of the biochemical processes involved in human sialic acid biology and the genes known to be associated with them.

Both the large number of genes involved in human sialic acid biology and their high sequence diversity have been hypothesized to be a result of ancient selective pressures from paleopathogens that infected hominins millions of years ago and left their signatures on genes involved in sialic acid biology (Varki 2009). The most striking example involves the *Alu*-mediated inactivation of the *CMAH* gene specifically in the human lineage (Varki 2001; Chou et al. 1998; Irie et al. 1998): the enzyme encoded by this gene is responsible for the conversion of N-acetylneuraminic acid (Neu5Ac) to N-glycolylneuraminic acid (Neu5Gc), and Neu5Ac is the main form of sialic acid in humans. This change in sialic acid composition has been suggested to act as a means to escape infection by *Plasmodium reichenowi*, a parasite that preferentially binds to Neu5Gc and is known to cause malaria in chimpanzees. Conversely, adoption of Neu5Ac likely made humans susceptible to infection by the malaria parasite *Plasmodium falciparum*, which instead binds to Neu5Ac (Martin et al. 2005; Varki & Gagneux 2009). Additionally, in contrast to humans, the inhibitory receptor Siglec-5 and its immunostimulatory counterpart Siglec-14 in great apes lack the essential arginine residue required for optimal binding with sialic acids, likely as a consequence of the above-mentioned switch from Neu5Gc to Neu5Ac in the human lineage (Angata et al. 2006). Finally, molecular evolutionary analyses in primates and rodents have shown that genes involved in sialic acid recognition show a considerable degree of sequence divergence, even between closely related species, suggestive of ancient positive selection in response to paleopathogens (Altheide et al. 2006).

The examples above suggest that sialic acid biology genes experienced adaptive changes early in the evolutionary radiation of primates, including hominids and hominins. However, whether these same genes also experienced positive selection during recent human evolution (i.e., in the last 250,000 years), reflective of adaptation to contemporary pathogens, remains an open question. To test this hypothesis, we estimated the population-level genetic variation of the protein-coding regions of sialic acid biology genes, and carried out tests aimed to detect hard selective sweeps, including tests of deviation from neutrality (measured by Tajima’s *D*; Tajima 1989), population differentiation (measured by *F*_*ST*_; Weir & Cockerham 1984), and extended haplotype homozygosity (measured by *nS*_*L*_; Ferrer-Admetlla et al. 2014) using the 1000 Genomes Project sequencing data from 1,668 humans belonging to three ethnic groups (Auton et al. 2015). To also test for the possible action of soft selective sweeps, we employed H12 (Garud et al. 2015), a measure of modified pooled haplotype homozygosity. To determine whether sialic acid biology genes exhibit patterns of genetic variation that deviate from neutral expectations, we compared the values of each of these four measures for sialic acid biology genes against those from sequences of the same length simulated under neutral evolution assuming a realistic model of human demographic history (Haller & Messer 2017; Gravel et al. 2011; Messer 2013). Examination of patterns of genetic variation of sialic acid biology genes showed that, irrespective of their functional category, most of them conform to neutral expectations. Sialic acid biology genes that deviate from neutrality exhibit evidence of either temporally or spatially varying positive selection. For example, both recognition genes *SIGLEC5* and *SIGLEC12* show *nS*_*L*_ values in Europeans that are consistent with the occurrence of a selective sweep approximately 20,000 years ago, whereas the Tajima’s *D* value of the recycling and degradation gene *SIAE* suggests that it experienced a hard selective sweep long before the out-of-Africa migration, approximately 250,000 years ago. Combined with previous work showing the widespread occurrence of ancient positive selection on genes involved in sialic acid biology prior to the emergence of modern humans (Altheide et al. 2006), our results suggest that the evolution of human sialic acid biology genes has been more strongly influenced by ancient primate pathogens rather than by contemporary cosmopolitan or geographically-restricted human pathogens.

## Methods

### Genotype Dataset

To examine whether sialic acid biology genes have experienced positive selection during recent human evolution, for all analyses we used the genotype data for 1,668 individuals from three ethnic backgrounds (661 Africans, 503 Europeans, and 504 East Asians) from Phase 3 of the 1000 Genomes Project (for information on data sources, see File S1) (Auton et al. 2015).

### Sialic Acid Biology Genes

To identify all the genes involved in human sialic acid biology, we started from a previously published list of 55 loci (Altheide et al. 2006) and added *SIGLEC14* and *SIGLEC16*, which were discovered more recently, as well as *SIGLEC15*, which was not included in the previous study. As sex-linked genes tend to exhibit different patterns of genetic variation than genes located on autosomal chromosomes (Schaffner 2004), we excluded the two genes that are located on the X chromosome, *L1CAM* and *RENBP*; in addition, we removed *CMAH*, the sole gene involved in modification of newly synthesized sialic acids, because it is a non-functional pseudogene in humans (Chou et al. 1998; Irie et al. 1998). Thus, 55 genes were retained for subsequent analyses (Figure 1).

To extract genotypes for these 55 genes, we used *VCFtools*, version 0.1.13 (Danecek et al. 2011) (for specific commands used, see File S1), with the gene coordinates (GRCh37p.13) obtained from Ensembl (release 75) via BioMart as input; indels were excluded from our analyses. Compressed VCF files and index files required for subsequent analyses were created using *tabix*, version 0.2.6 (Li 2011).

### Measuring signatures of recent positive selection in human sialic acid biology genes

To determine the levels of genetic polymorphism of sialic acid biology genes, we first calculated pairwise nucleotide diversity (π) (Nei & Li 1979) using *PopGenome*, version 2.1.6 (Pfeifer et al. 2014). As there may have been variation in when and where human sialic acid biology genes experienced positive selection, we next used four different measures aimed to capture genetic signatures left from the action of selection at different time ranges during recent human evolution (Figure 2) (Sabeti et al. 2006).

**Figure 2.**
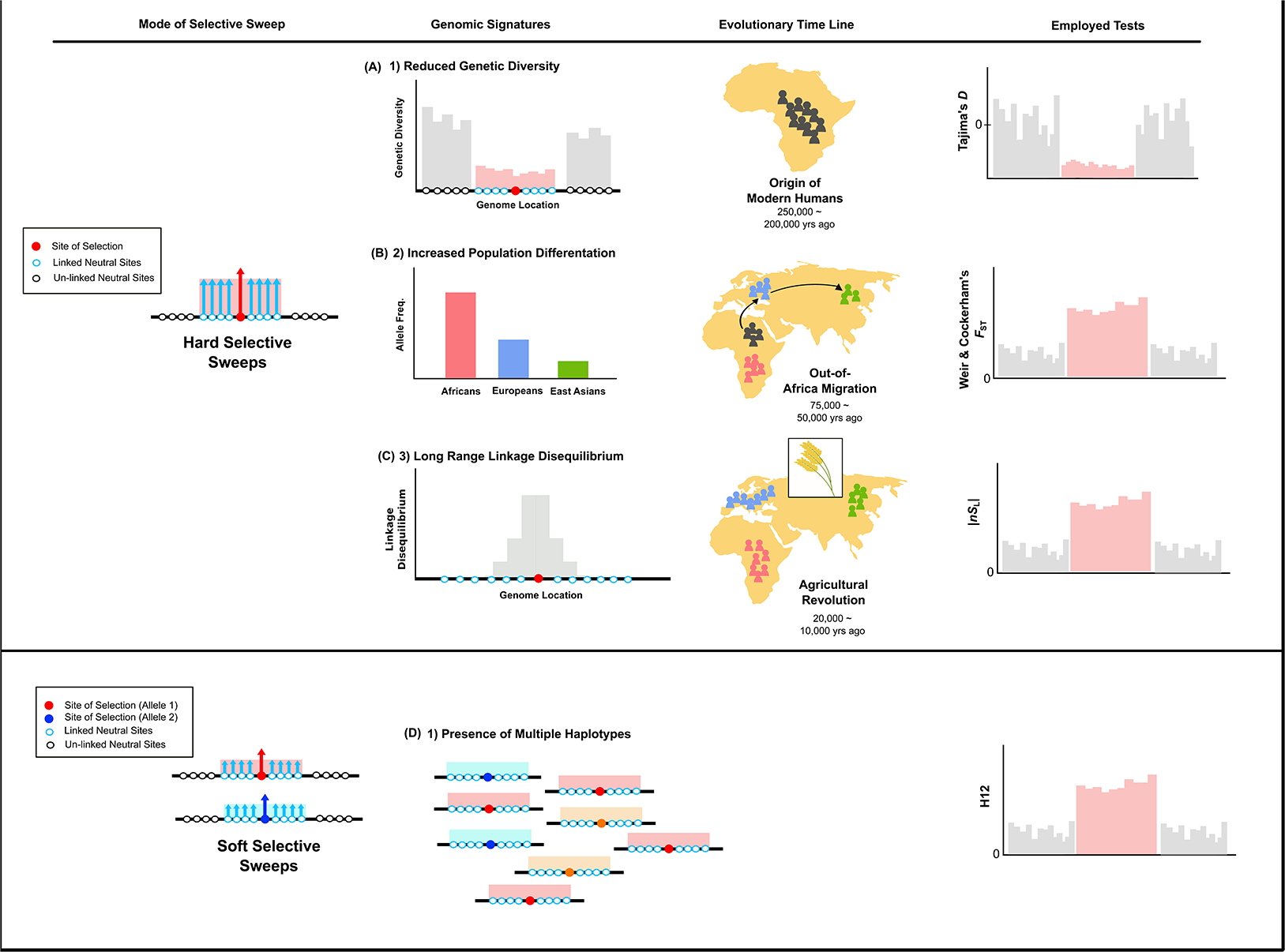
Summary of the population genetics measures used in this study. This figure depicts the nature of the genomic signatures left by varying modes of selective sweeps occurring at different evolutionary time points, the measures that we use to detect such signatures, and the expected values for regions that have experienced selective sweeps (depicted in pink) compared to regions that have not (depicted in gray). (A) In a hard selective sweep model, a single, advantageous new variant rises to high frequency (‘sweeps’) within the population, dramatically reducing variation in the nearby (linked) neutral variants. This results in a signature of reduced levels of genetic diversity in the genomic region surrounding the selected variant. The Tajima’s *D* neutrality index assesses the extent of the abundance of rare variants resulting from such selective sweep events. As mutagenesis is rare, this signature persists for a relatively long period of time, and has been proposed to detect selective events that happened around the origin of modern humans approximately 250,000 years ago. (B) Approximately around 75,000 - 50,000 years ago, modern humans migrated out of Africa to inhabit different parts of the world, resulting in exposure to new selective pressures (e.g., pathogens). Subsequent local adaptation events would have resulted in differences in allele frequencies among distinct populations (i.e. population differentiation), which can be quantified by Weir & Cockerham’s *F*_ST_ index. (C) Another genomic signature of hard selective sweeps is the presence of one dominant extended haplotype within the population, which stems from the hitch-hiking of linked neutral variants alongside the beneficial variant. As recombination events eventually break down the haplotype structure, this signature persists for a relatively short period of time (i.e. 20,000 - years). This signature can be quantified by measures of extended haplotype homozygosity such as *nS*_L._ (D) In contrast to the hard selective sweeps, soft selective sweeps occur when standing variation or multiple *de novo* mutations introduce several beneficial alleles into the population, resulting in an increase in the frequency of more than one, independent haplotypes. Measures of pooled haplotype homozygosity that combine the frequencies of the most common and second-most-common haplotypes (i.e. H12) have been shown to possess increased power to capture the signatures resulting from such soft selective sweeps.

To detect signatures left by hard selective sweeps that occurred approximately 250,000 years ago, we calculated Tajima’s *D* (Figure 2A) (Tajima 1989) using the R package *PopGenome*, version 2.1.6 for each of the 55 genic regions (i.e. regions defined by the gene coordinates obtained via Ensembl) of the sialic acid biology genes (Pfeifer et al. 2014). To detect local selection that occurred after the out-of-Africa migration approximately 75,000 years ago, we calculated both weighted and mean Weir & Cockerham’s fixation index (*F*_ST_) (Figure 2B) (Weir & Cockerham 1984) between all three ethnic groups for the genic regions of sialic acid biology genes using *VCFtools* (Danecek et al. 2011). To capture signatures of selective sweeps that occurred approximately 20,000 years ago (Sabeti et al. 2006; Voight et al. 2007), we calculated the *nS*_L_ statistic (number of segregating sites by length; Ferrer-Admetlla et al. 2014) (Figure 2C) (Sabeti et al. 2006) using *Selscan*, version 1.2.0 (Szpiech & Hernandez 2014). For each gene, we calculated *nS*_L_ for the region extending 100kb upstream and downstream using the default settings of *Selscan*; the only deviation from the default settings was that we used 0.01, instead of 0.05, for the minor allele frequency (MAF) cut-off value. As extended haplotype methods are thought to detect selection events that happened after the out-of-Africa migration, we calculated this metric for each of the three ethnic groups, rather than globally. We used the maximum absolute *nS*_L_ values calculated over the entire window to represent each sialic acid biology gene and report un-standardized *nS*_L_ values in this paper.

Tajima’s *D*, Weir & Cockerham’s *F*_ST_, and the *nS*_L_ statistic are designed to capture signatures of completed or ongoing hard selective sweeps; therefore, they have limited power to detect incidences of soft selective sweeps, in which standing variation or multiple *de novo* mutations introduce several beneficial alleles into a population (Pennings & Hermisson 2006a; Messer & Petrov 2013). To detect soft sweeps, we used the H12 index (Figure 2D) (Garud et al. 2015), a modified haplotype homozygosity index that combines the frequency of the most common and second most common haplotypes in a given sample (Pennings & Hermisson 2006b). We modified the original python script written by Garud to carry out the calculations of H12 (Garud et al. 2015) on the genic regions of sialome genes.

### Simulations of Neutral Evolution

The distribution of each of the four metrics in the absence of selection can be influenced by demographic events and random processes, such as genetic drift. Therefore, to assess the probability that observed values in sialic acid biology genes reflect the action of selection, we compared them to those “expected” from a realistic neutral model that accounts for human demography. To identify any sialic acid biology genes that significantly deviate from selective neutrality in any of four measures, we conducted simulations of neutral evolution using *SLiM*, version 2.4.1 (Haller & Messer 2017; Messer 2013). To account for the confounding effects of past demographic events, we incorporated previously calculated demographic parameters (Gravel et al. 2011), which combined low-coverage whole genome data and high-coverage targeted exon data of the pilot phase 1000 Genomes Project and estimated the demographic parameters for the three ethnic groups in our study. More specifically, the simulation model assumes that: a) approximately 148,000 years ago, the ancestral African population experienced a population expansion from an initial effective population size of 7,310 to 14,474, b) the out-of-Africa migration occurred approximately 51,000 years ago, c) the subsequent Eurasian split happened approximately 23,000 years ago, d) after the Eurasian split, a bottleneck event in the European population reduced its effective size to 1,032 individuals, e) for the last 23,000 years, both the European and the East Asian populations each experienced an exponential growth in effective population size (exponential coefficients for Europeans: 0.0038; exponential coefficients for East Asians: 0.0048), and f) both the recombination rate (1.0 x10^−8^ recombination events per bp) and the mutation rate (2.36 x 10^−8^ mutations per bp) were fixed. More detailed information on the parameters used for the simulations can be found in the Supplementary File S1.

Using this model and parameters, we simulated the genotypes of 661, 503, and 504 individuals corresponding to the number of individuals that we have genotype data from the African, European, and East Asian groups, respectively. For each gene, we carried out 5,000 simulations to create neutrally-evolved sequences of the same length. We next used these simulated sequences to calculate Tajima’s *D*, Weir & Cockerham’s *F*_ST_, and H12 values as described above. To calculate *nS*_L_ values, we used *SLiM* to simulate 2,500 genomic regions that extend 100kb upstream and downstream a “simulated sialic acid biology gene”. *nS*_L_ values were calculated on these regions using *Selscan* as described above. For all tests, we calculated the *p*-value for a given gene as the proportion of values on simulated sequences that were equal to or more extreme than the observed value. We used a *p*-value of 0.05 as cutoff for significance: a gene associated with a *p*-value lower than 0.05 likely indicates 1) deviation from neutrality, and 2) patterns of variation are due to the action of positive selection and not due to demographic events.

### Statistical Analysis

To compare the patterns of genetic variation across different functional categories of sialic acid biology genes, we conducted pairwise Mann-Whitney *U* (MW*U*) tests (two-sided): the exact *p*-values were calculated for each comparison via the *coin* package, version 1.1-3, in the R programming environment (Hothorn et al. 2008). *P*-values calculated for comparisons among functional categories were adjusted *post hoc* to correct for the testing of multiple hypotheses via the Bonferroni method using R.

### Data Availability

**File S1 contains detailed descriptions of the methods used and the locations of the data files.**

**File S2 lists the genomic coordinates of the 55 sialic acid biology genes.**

## Results

### Most sialic acid biology genes do not exhibit signatures of recent positive selection

To test the hypothesis that human sialic acid biology genes experienced recent positive selection resulting from hard selective sweeps approximately 250,000 years ago (i.e. around the origin of modern humans), we calculated Tajima’s *D* for each of the 55 sialic acid biology genes across our sample of 1,668 individuals (Figure 2A & Table S1). Comparison of each gene’s Tajima’s *D* value against the distribution of Tajima’s *D* values estimated from 5,000 simulated sequences under neutrality showed that only 2 / 55 sialic acid biology genes exhibited significantly extreme values (*SIAE*, involved in recycling, degradation functions: *p*-value = 0.024; and *ST3GAL2*, involved in activation, transport, and transfer: *p*-value *=* 0.003; Figure 3 & Table S1).

**Figure 3.**
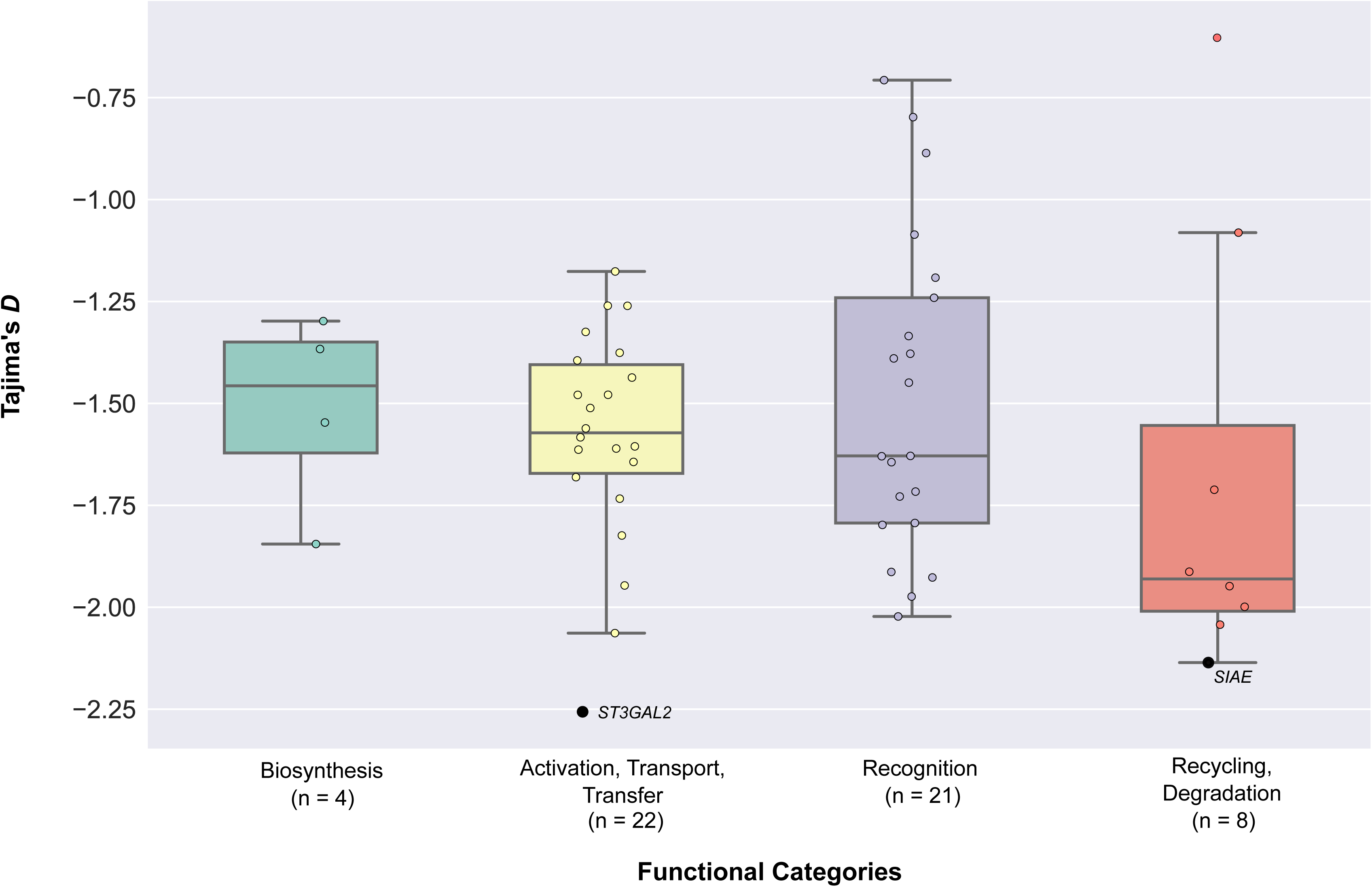
The distribution of Tajima’s *D* values across the four functional categories of sialic acid biology genes. The graph presents the values of Tajima’s *D* calculated for all 1,668 individuals of the 1000 Genomes Project. The 55 sialic acid biology genes are displayed according to the functional categories they belong to. Genes exhibiting significant deviation from neutral expectations (i.e. genes with *p*-values less than 0.05) are highlighted in black and their names are shown next to the data points; the names of all other (non-significant) genes have been omitted. The values of Tajima’s *D* for each of the 55 genes can be found in Table S1. The number of genes belonging to each category is shown below the name of each functional category.

To test the hypothesis that human sialic acid biology genes experienced recent local positive selection after the major human migration out of Africa approximately 75,000 years ago, we calculated Weir & Cockerham’s *F*_ST_ between our three human populations for each of the sialic acid biology genes (Figure 2B & Table S1). We found that 53 / 55 sialic acid biology genes did not exhibit significantly higher levels of population differentiation compared to the values calculated for 5,000 neutrally simulated sequences (Figure 4 & Table S1); the two exceptions were the biosynthesis-involved *NANP* (*p*-value = 0.045) and the recognition-involved *SELP* (*p*-value = 0.035).

**Figure 4.**
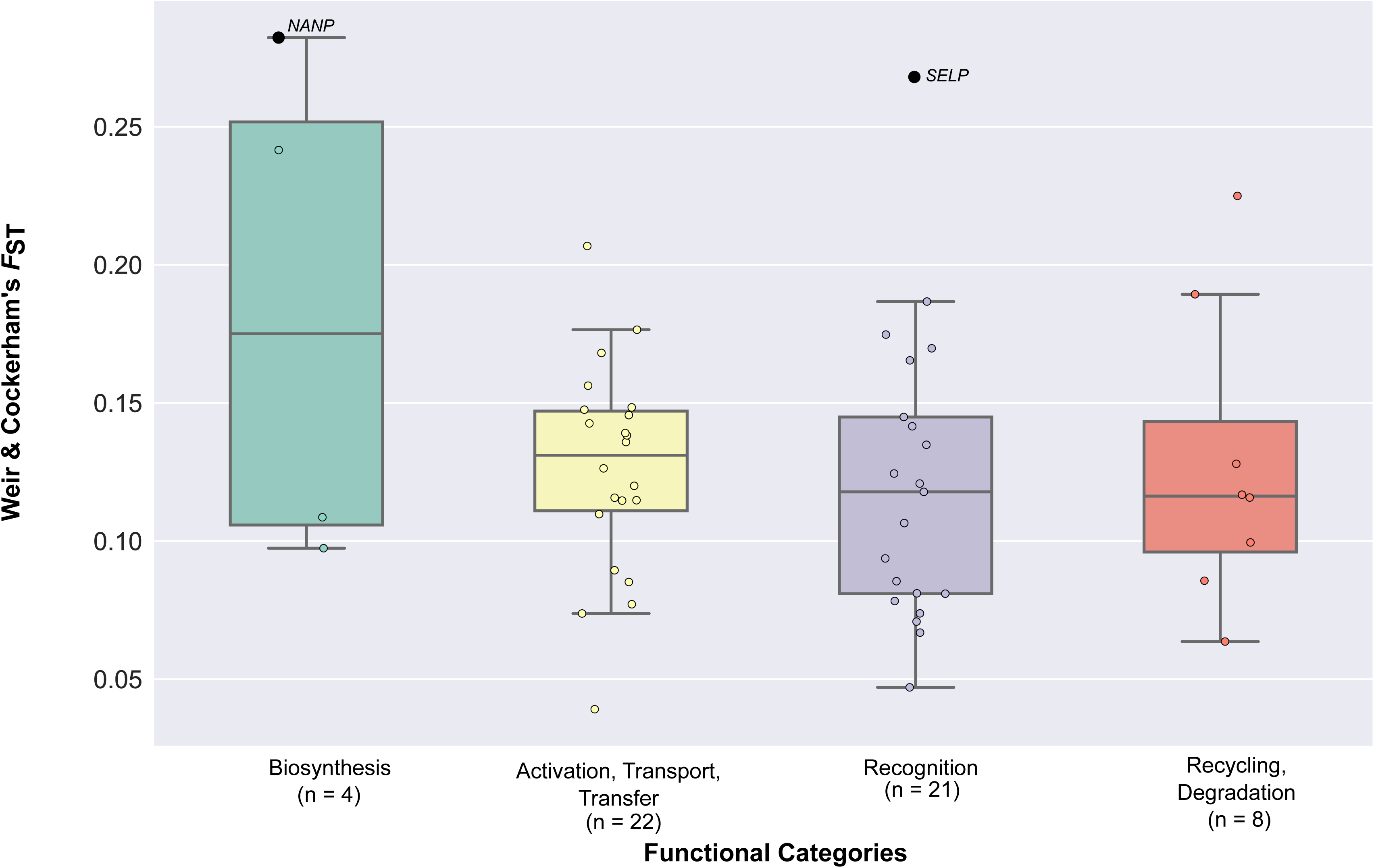
The distribution of Weir & Cockerham’s *F*_ST_ values across the four functional categories of sialic acid biology genes. The graph presents the values of weighted Weir & Cockerham’s *F*_ST_ values calculated by pairwise comparisons of the three ethnic groups. The 55 sialic acid biology genes are displayed according to the functional categories they belong to. Genes exhibiting significant deviation from neutral expectations (i.e. genes with *p*-values less than 0.05) are highlighted in black, their names shown next to the data points; the names of all other (non-significant) genes have been omitted. The values of the *F*_ST_ for each gene, along with the values of the mean Weir & Cockerham’s *F*_ST_, can be found in Table S1. The number of genes belonging to each category is shown below the name of each functional category.

To test the hypothesis that human sialic acid biology genes experienced selective sweeps in one or more human populations approximately 20,000 years ago (i.e. around or shortly prior to the Agricultural Revolution), we calculated *nS*_L_ values for the regions flanking 100kb up- and down-stream of sialic acid biology genes within each ethic group (Figure 2C & Table S2). For all three ethnic groups, we found that most sialic acid biology genes did not deviate from neutral expectations: only five genes (four involved in recognition and one in recycling, degradation) in Europeans (*SIGLEC5*: *p*-value = 0.0004; *SIGLEC6*: *p*-value = 0.0004; *SIGLEC12*: *p*-value = 0.0004; *SIGLEC14*: *p*-value = 0.0004; and *NEU2*: *p*-value = 0.036; Figure 5B & Table S2), five genes (three involved in recognition and two in activation, transport, and transfer) in Africans (*LAMA2*: *p*-value = 0.018; *ST6GALNAC1*: *p*-value = 0.002; *ST6GALNAC2*: *p*-value = 0.001; *SIGLEC8*: *p*-value = 0.002; and *SIGLEC10*: *p*-value = 0.002; Figure 5A & Table S2), and four genes (two involved in recognition and two in activation, transport, and transfer) in East Asians (*CD22*: *p*-value = 0.046; *MAG*: *p*-value = 0.046, *ST6GAL1*: *p*-value = 0.001; and *ST6GALNAC5*: *p*-value = 0.030; Figure 5C & Table S2) exhibited significant *p*-values. Notably, the genes exhibiting significant deviations from neutral expectations were different for each ethnic group.

**Figure 5.**
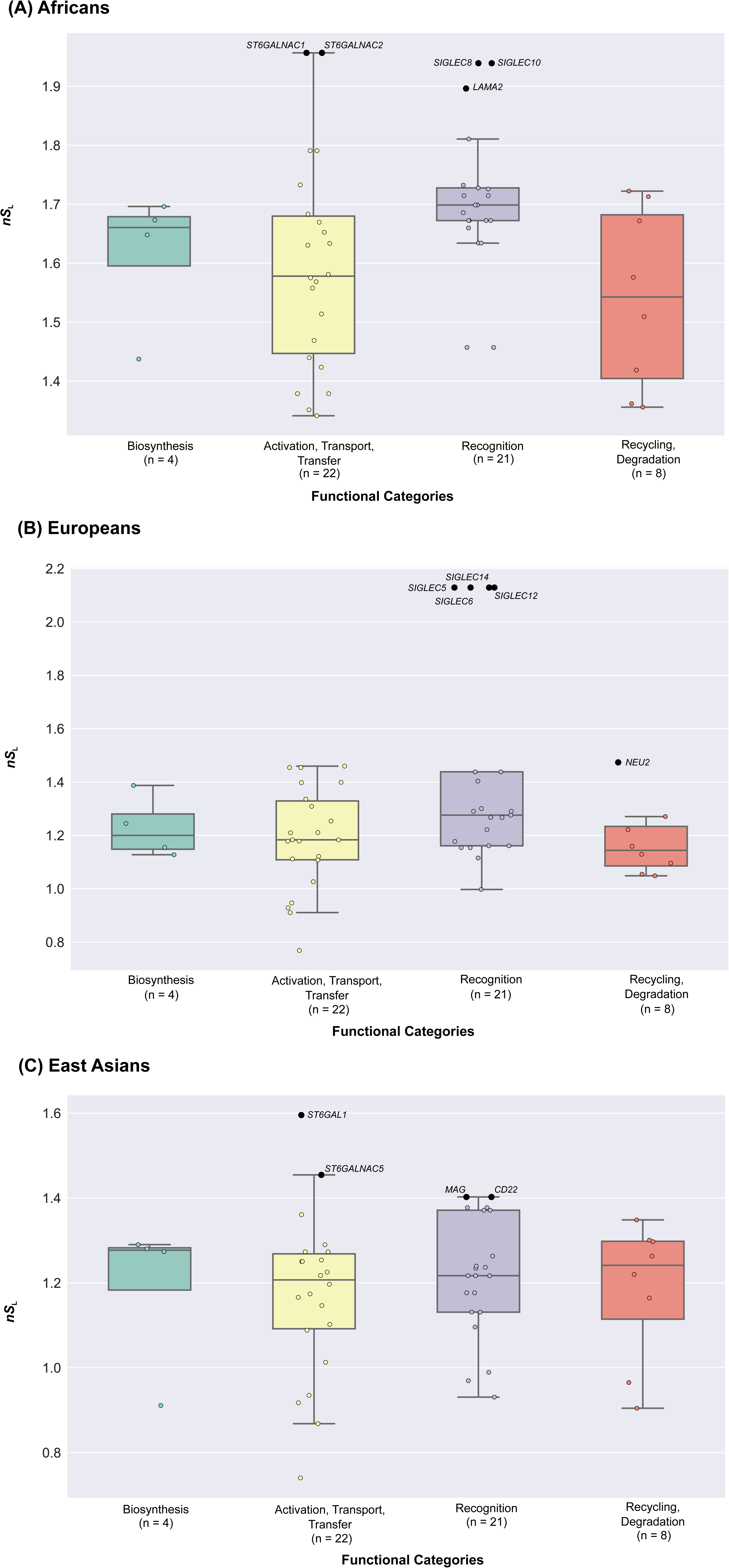
The distribution of *nS*_L_ values across the four functional categories of sialic acid biology genes. The graph presents the values of *nS*_L_ values calculated for each ethnic group (Africans: (A), Europeans: (B), East Asians: (C)). Unstandardized *nS*_L_ values were calculated in windows that expand 100kb up and downstream of a given sialic acid biology gene: the maximum value of all the absolute *nS*_L_ values in a given window was used to represent each gene. The 55 sialic acid biology genes are displayed according to the functional categories they belong to. Genes exhibiting significant deviation from neutral expectations (i.e. genes with *p*-values less than 0.05) are highlighted in black, their names shown next to the data points; the names of all other (nonsignificant) genes have been omitted. The values of the *nS*_L_ for each gene can be found in Table S2. The number of genes belonging to each category is shown below the name of each functional category.

Finally, to test the hypothesis that the sialic acid biology genes experienced soft selective sweeps, we calculated the modified pooled haplotype homozygosity index H12 for each of the 55 sialic acid biology genes (Figure 2D; Table S1). 10 / 55 sialic acid biology genes showed H12 values that significantly deviated from neutral expectations (*NANS*: *p*-value = 0.0004; *NEU1*: *p*-value = 0.027; *NEU3*: *p*-value = 0.001; *NPL*: *p*-value = 0.0004; *SIAE*: *p*-value = 0.001; *SIGLEC6*: *p*-value = 0.023; *SLC35A1*: *p*-value = 0.001; *ST3GAL5*: *p*-value = 0.002; *ST6GALNAC1*: *p*-value = 0.032; *ST8SIA4*: *p*-value = 0.015; Figure 6 & Table S1). Of these 10 genes, four function in sialic acid activation, transport, and transfer (*SLC35A1*, *ST3GAL5*, *ST6GALNAC1*, and *ST8SIA4*), four in recycling and degradation (*NEU1*, *NEU3*, *NPL*, and *SIAE*), one in recognition (*SIGLEC6*) and one in biosynthesis (*NANS*).

**Figure 6.**
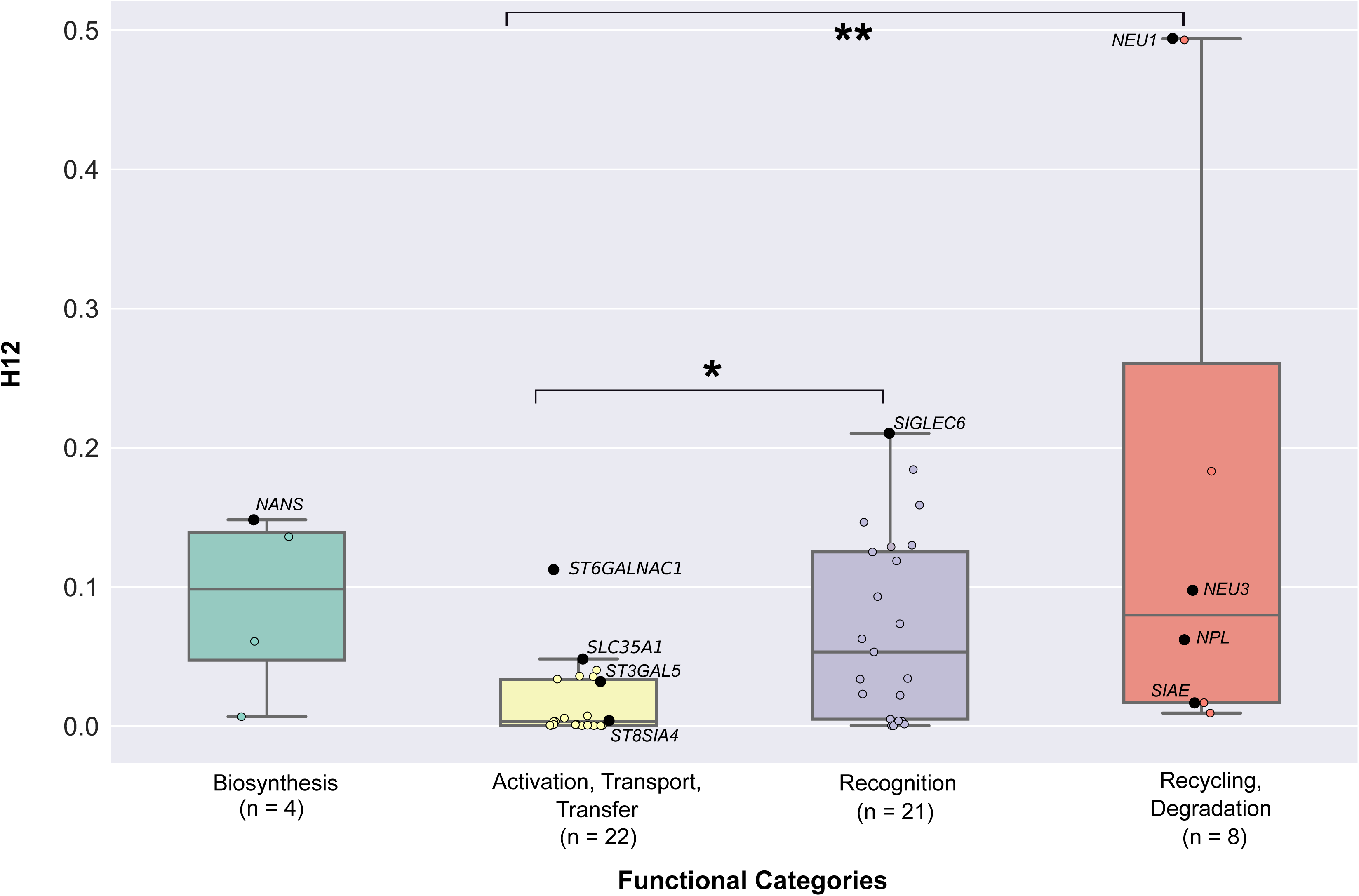
The distribution of H12 values across the four functional categories of sialic acid biology genes. The graph presents the values of H12 values calculated across all three ethnics groups. The 55 sialic acid biology genes are displayed according to the functional categories they belong to. Significant pairwise functional comparisons are indicated with asterisks (*: *p*-value < 0.05; **: *p*-value < 0.01). Genes exhibiting significant deviation from neutral expectations (i.e. genes with *p*-values less than 0.05) are highlighted in black, their names shown next to the data points; the names of all other (non-significant) genes have been omitted. The values of the H12 for each gene can be found in Table S1. The number of genes belonging to each category is shown below the name of each functional category.

### The four functional categories of sialic acid biology genes do not significantly differ in their signatures of recent positive selection

We next examined whether the four distinct functional categories of sialic acid biology genes (Figure 1) exhibit statistically significant differences in their extent of polymorphism, allele frequency spectra, extent of population differentiation, and haplotype homozygosity by comparing the patterns of nucleotide diversity (π), Tajima’s *D*, *F*_ST_, *nS*_L_, and H12 values across the four functional categories. 4 / 5 measures (π: Figure 7 & Table S3A; Tajima’s *D*: Figure 3 & Table S3B; *F*_ST_: Figure 4 & Table S3C; *nS*_L_: Figure 5A-C & Table S3D-F) did not show significant differences among the four functional categories. In contrast, comparison of H12 values across the four functional categories showed significant differences between activation, transport, transfer and recognition (*U*-value = 114, adjusted *p*-value = 0.023; Figure 6 & Table S3G) and between activation, transport, transfer and recycling, degradation (*U*-value = 23, adjusted *p*-value = 0.008; Figure 6 & Table S3G).

**Figure 7.**
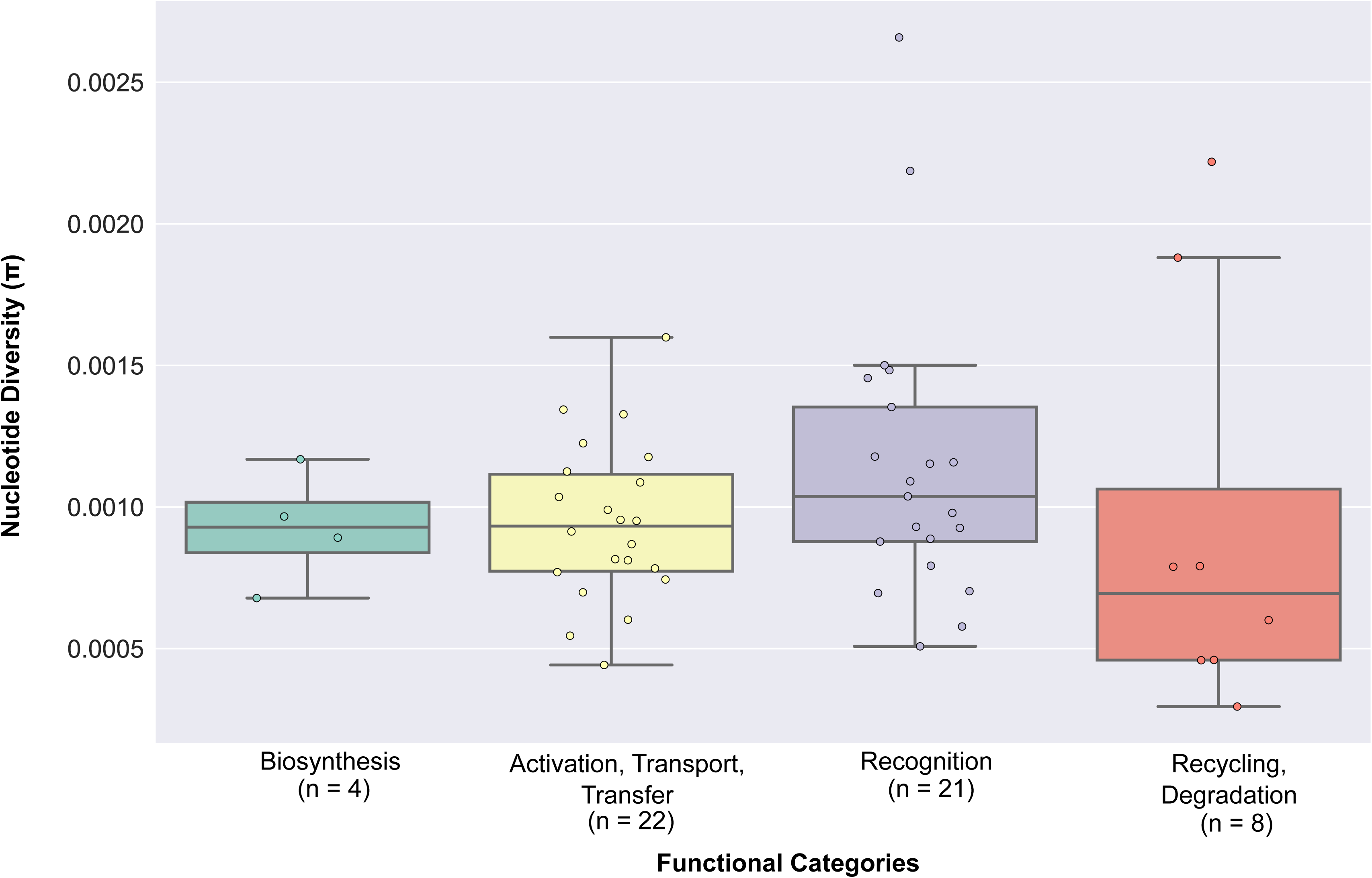
The distribution of nucleotide diversity values across the four functional categories of sialic acid biology genes. The graph presents the nucleotide diversity values calculated across all three ethnic groups. The 55 sialic acid biology genes are displayed according to the functional categories they belong to. The nucleotide diversity value for each gene can be found in Table S1. The number of genes belonging to each category is shown below the name of each functional category.

## Discussion

In this study, we tested the hypothesis that loci involved in human sialic acid biology experienced recent positive selection by employing four population genetic tests designed to detect selective signatures at different time points during the evolution of modern humans. In all of our analyses, the majority of sialic acid biology genes did not significantly deviate from neutral expectations.

One likely explanation of our results is that, for the most part, genes involved in sialic acid biology have not been targets of positive selection in the last 250,000 years of human evolution. Previous studies have shown that sialic acid biology genes were subject to strong ancient selective pressures that have resulted in species-specific adaptations (Varki 2001; Angata et al. 2004; Altheide et al. 2006; Varki & Gagneux 2009), suggesting that they likely evolved in response to ancient pathogens. The function of sialic acid biology genes, i.e., the generation of host-specific sialic acids and recognition of those self-associated molecular patterns (SAMPs) by cognate receptors, contributes to the detection of missing self (Medzhitov & Janeway 2002). It is possible that evolution of such markers of normal self, and the ability to recognize them correctly, have been influenced by ancient, but not, recent pathogens. For instance, the human-specific *Alu*-mediated inactivation of *CMAH*, the protein product of which is responsible for generating Neu5Gc from its precursor Neu5Ac, has been suggested as a means of escaping infection by *Plasmodium reichenowi*, a malaria-causing parasite that infects other great apes (Martin et al. 2005; Varki & Gagneux 2009). In addition, human-specific mutation of an essential arginine residue required for sialic acid binding in Siglec-12 has been suggested to be evolutionarily related to the loss of Neu5Gc in humans (Varki 2009). Several other human-specific changes in sialic acid biology (e.g., deletion of *SIGLEC13* in humans or increased expression of *SIGLEC1* on human macrophages compared to other primates) could have resulted from selective pressures exerted by paleopathogens (Varki 2009; Wang et al. 2012). It should be noted that several other innate-immunity genes have been shown to lack any signatures of positive selection in more recent evolutionary time scales (Mukherjee et al. 2009; Siddle & Quintana-Murci 2014).

An alternative, not mutually exclusive, possibility is that sialic acid biology genes may have experienced recent positive selection but – as the strength of selection can fluctuate over time and space – any signatures left from such selective episodes was not strong enough to alter the allele frequency spectra and extent of haplotype homozygosity of sialic acid biology genes within a single population or across multiple populations to detectable levels (Bell 2010; Pritchard et al. 2010). For example, even our most sensitive measure (the H12 index, which can discern past soft selective sweeps) is still biased towards detecting hard selective sweeps and loses power when sweeps are too soft (i.e. when selection has acted on numerous variants, resulting in a high number of haplotypes being present in the population) (Garud et al. 2015). Therefore, it is possible that signatures of selection left by very soft selective sweeps are not being detected by the methods employed in this study. In addition, host defense is a complex biological process that involves interactions between numerous genes belonging to diverse biological pathways. As such, it is feasible that selection has operated not on individual sialic acid biology genes, but on the set of pathways that comprise sialic acid biology, resulting in only subtle changes in allele frequency spectra and haplotype homozygosity patterns of individual genes that could not be detected by the gene-specific tests employed in our study. The occurrence of such polygenic adaptation (Berg & Coup 2014; Daub et al. 2013; Daub et al. 2015) could similarly result in non-significant deviations from neutral expectations at the level of individual loci.

Among the sialic acid biology genes that did show evidence of recent positive selection, most were recovered by the haplotype-based H12 index (10 genes), with many fewer recovered by either the haplotype-based *nS_L_* (4-5 per population) or the two frequency spectra-based metrics (Tajima’s *D* and *F_ST_*; 2 each) (Figure 8). The key difference among these metrics is that H12 has increased power to detect soft selective sweeps whereas the other three are designed to detect hard selective sweeps, suggesting that sialic acid biology genes likely experienced more soft than hard sweeps. This is consistent with a previous study suggesting that hard selective sweeps were likely rare during modern human evolution (Hernandez et al. 2011). Interestingly, genes that underwent hard selective sweeps were often involved in recognition of sialic acids, whereas genes that underwent soft sweeps were more often in the ‘activation, transport, transfer’ and ‘recycling, degradation’ categories (Figures 2 and 8).

**Figure 8.**
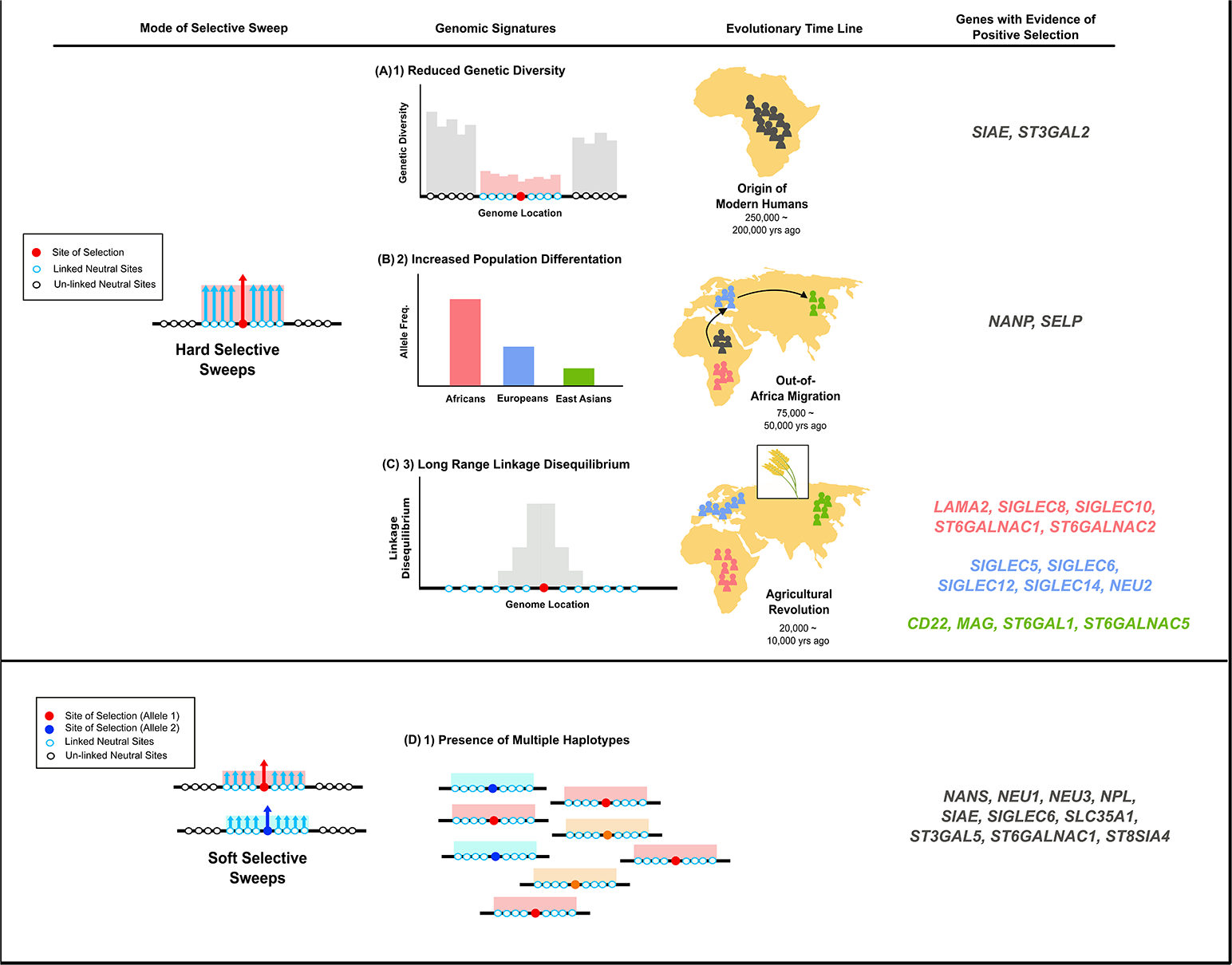
Few sialic acid biology genes have experienced both temporally and spatially varying modes of selective sweeps in recent human history.

We also found that the identity of the genes exhibiting significant deviations from neutral expectations was different among the various tests of selection employed in this study, and, in the case of *nS*_L_, among the three ethnic groups studied. The only overlap was for *SIGLEC6*, which was shown to exhibit significant values of both *nS*_L_ calculated in Europeans and H12. The lack of overlap likely reflects the differences in sensitivities of the four metrics (see preceding paragraph), and hence the occurrence of spatially and/or temporally varying selective events. For instance, it has been demonstrated that tests that quantify the extent of excess or depletion of rare alleles (e.g., Tajima’s *D*) have the most power to detect hard selective sweeps that happened at earlier time points, since the low rate of mutagenesis in humans does not erase such signatures (Sabeti et al. 2006). Similarly, *nS*_L_ has been shown to exhibit high power to detect signatures of selective sweeps that happened more recently and have resulted in only incomplete or partial fixation of alleles (Sabeti et al. 2006; Ferrer-Admetalla et al. 2014; Voight et al. 2007).

In summary, considered jointly with previous evidence on the widespread occurrence of ancient positive selection on genes involved in sialic acid biology in hominids and hominins (Altheide et al. 2006; Angata et al. 2006; Hayakawa & Varki 2011; Varki & Gagneux 2009), these results support the hypothesis that human sialic acid biology genes bear many and strong evolutionary signatures of adaptation in response to ancient primate pathogens, but rather few and weak adaptive signatures in response to contemporary cosmopolitan or geographically-restricted human pathogens. Investigation of the possible actions of alternative modes of selection (e.g., balancing selection; Charlesworth 2006; Gao et al. 2015) or extremely recent selection (e.g., the singleton density score designed to detect selective events 3,000 – 2,000 years ago; Field et al. 2016) in these, as well as in additional human populations, will lead to a more thorough understanding of the evolution of sialic acid biology during modern human history.

## Acknowledgements

This work was conducted in part using the resources of the Advanced Computing Center for Research and Education at Vanderbilt University. MM was partially supported by the Graduate Program in Biological Sciences at Vanderbilt University. This work was supported in part by the March of Dimes through the March of Dimes Prematurity Research Center Ohio Collaborative, a Burroughs Wellcome Fund Preterm Birth Initiative Award (to J.A.C. and A.R.), and by the National Science Foundation (DEB-1442113 to A.R.).

## References

Altheide TK et al. 2006. System-wide Genomic and Biochemical Comparisons of Sialic Acid Biology Among Primates and Rodents: EVIDENCE FOR TWO MODES OF RAPID EVOLUTION. J Biol Chem. 281:25689–25702.

Angata T, Hayakawa T, Yamanaka M, Varki A, Nakamura M. 2006. Discovery of Siglec-14, a novel sialic acid receptor undergoing concerted evolution with Siglec-5 in primates. FASEB J. 20: 1964–1973.

Angata T, Margulies EH, Green ED, Varki A. 2004. Large-scale sequencing of the CD33-related Siglec gene cluster in five mammalian species reveals rapid evolution by multiple mechanisms. PNAS. 101: 13251–13256.

Angata T, Varki A. 2002. Chemical Diversity in the Sialic Acids and Related α-Keto Acids: □ An Evolutionary Perspective. Chem. Rev. 102: 439–470.

Bell G. 2010. Fluctuating selection: the perpetual renewal of adaptation in variable environments. Philos. Trans. R. Soc. Lond. B. Biol. Sci. 365: 87–97.

Auton A et al. 2015. A global reference for human genetic variation. Nature. 526: 68–74.

Berg JJ, Coop G. 2014. A Population Genetic Signal of Polygenic Adaptation. PLoS Genet. 10(8): e1004412.

Charlesworth D. 2006. Balancing Selection and Its Effects on Sequences in Nearby Genome Regions. PLoS Genet. 2(4): 364.

Chou HH et al. 1998. A mutation in human CMP-sialic acid hydroxylase occurred after the Homo-Pan divergence. Proc. Natl. Acad. Sci. USA. 95: 11751–11756.

Cohen M, Varki A. 2010. The Sialome–Far More Than the Sum of Its Parts. OMICS. 14: 455–464.

Danecek P et al. 2011. The variant call format and VCFtools. Bioinformatics. 27: 2156–2158.

Daub JT et al. 2013. Evidence for polygenic adaptation to pathogens in the human genome. Mol. Biol. Evol. 30: 1544–1558.

Daub JT, Dupanloup I, Robinson-Rechavi M, Excoffier L. 2015. Inference of Evolutionary Forces Acting on Human Biological Pathways. Genome Biol. Evol. 7: 1546–1558.

Ferrer-Admetalla A, Liang M, Korneliussen, Nielsen R. 2014. On Detecting Incomplete Soft or Hard Selective Sweeps Using Haplotype Structure. Mol. Biol. Evol. 31: 1275–1291.

Field Y et al. 2016. Detection of human adaptation during the past 2000 years. Science. 354: 760–764.

Gao Z. Przeworski M. Sella G. 2015. Footprints of ancient-balanced polymorphisms in genetic variation data from closely related species. Evolution. 69: 431–446.

Garud NR, Messer PW, Buzbas EO, Petrov DA. 2015. Recent Selective Sweeps in North American *Drosophila melanogaster* Show Signatures of Soft Selective. PLoS Genet. 11(2): e1005004.

Gravel S et al. 2011. Demographic history and rare allele sharing among human populations. PNAS. 108: 11983–11988.

Haller BC, Messer PW. 2017. SLiM 2: Flexible, interactive forward genetic simulations. Mol. Biol. Evol. 34: 230–240

Hayakawa T, Varki A. 2011. Human-Specific Changes in Sialic Acid Biology in Primatology Monographs, pp.123–148 in Post-Genome Biology of Primates, edited by H. Hirai et al. Springer.

Hernandez RD et al. 2011. Classic Selective Sweeps Were Rare in Recent Human Evolution. Science. 331: 920–924.

Hothorn T, Hornik K, van de Wiel MA, Zeileis A. 2008. Implementing a Class of Permutation Tests: The coin Package. Journal of Statistical Software; 28: 1–23.

Irie A, Koyama S, Kozutsumi Y, Kawasaki T, Suzuki A. 1998. The Molecular Basis for the Absence of N-Glycolylneuraminic Acid in Humans. J Biol Chem. 273: 15866–15871.

Kelm S, Schauer R. 1997. Sialic Acids in Molecular and Cellular Interactions. Int Rev of Cytol 175: 137–240.

Li H. 2011. Tabix: fast retrieval of sequence features from generic TAB-delimited files. Bioinformatics. 27: 718–719.

Martin MJ, Rayner JC, Gagneux P, Barnwell JW, Varki A. 2005. Evolution of human-chimpanzee differences in malaria susceptibility: Relationship to human genetic loss of *N*-glycolylneuraminic acid. Proc. Natl. Acad. Sci. USA. 102: 12819–12824.

Medzhitov R, Janeway CA. 2002. Decoding the Patterns of Self and Nonself by the Innate Immune System. Science. 296: 298–300.

Messer PW. 2013. SLiM: Simulating Evolution with Selection and Linkage. Genetics. 194: 1037–1039.

Messer PW, Petrov DA. 2013. Population genomics of rapid adaptation by soft selective sweeps. Trends in Ecology & Evolution. 28: 1–11.

Mukherjee S, Sarkar-Roy N, Wagener DK, Majumder PP. 2009. Signatures of natural selection are not uniform across genes of innate immune system, but purifying selection is the dominant signature. Proc. Natl. Acad. Sci. USA. 106: 7073–7078.

Nei M, Li WH. 1979. Mathematical model for studying genetic variation in terms of restriction endonucleases. Proc. Natl. Acad. Sci. USA. 76: 5269–5273.

Pennings PS, Hermisson J. 2006a. Soft Sweeps II--Molecular Population Genetics of Adaptation from Recurrent Mutation or Migration. Mol Biol Evol. 23: 1076–1084.

Pennings PS, Hermisson J. 2006b. Soft Sweeps III: The Signature of Positive Selection from Recurrent Mutation. PLoS Genet. 2: e186.

Pfeifer B, Wittelsbϋrger U, Ramos-Onsins SE, Lercher MJ. 2014. PopGenome: An Efficient Swiss Army Knife for Population Genomic Analyses in R. Mol Biol Evol. 31: 1929–1936.

Pilatte Y, Bignon J, Lambré CR. 1993. Sialic acids as important molecules in the regulation of the immune system: pathophysiological implications of sialidases in immunity. Glycobiology. 3: 201–218.

Pritchard JK, Pickrell JK, Coop G. 2010. The Genetics of Human Adaptation: Hard Sweeps, Soft Sweeps, and Polygenic Adaptation. Current Biology. 20: R208–R215.

Sabeti PC et al. 2006. Positive Natural Selection in the Human Lineage. Science. 312: 1614–1620.

Schaffner SF. 2004. The X chromosome in population genetics. Nature Rev Genet. 5: 43–51.

Schauer R. 1982. Chemistry, Metabolism, and Biological Functions of Sialic Acids. Adv. Carbohydr. Chem. Biochem. 40: 31–234.

Siddle KJ, Quintana-Murci L. 2014. The Red Queen’s long race: human adaptation to pathogen pressure. Curr. Opin. Genet. Dev. 29: 31–38.

Szpiech ZA, Hernandez RD, 2014. selscan: An Efficient Multithreaded Program to Perform EHH-Based Scans for Positive Selection. Mol. Biol. Evol. 31: 2824–2827

Tajima F. 1989. Statistical method for testing the neutral mutation hypothesis by DNA polymorphism. Genetics. 123: 585–595.

Varki A. 2007. Glycan-based interactions involving vertebrate sialic-acid-recognizing proteins. Nature. 446: 1023–1029.

Varki A. 2001. Loss of N-glycolylneuraminic acid in humans: Mechanisms, consequences, and implications for hominid evolution. Am. J. Phys. Anthropol. 116: 54–69.

Varki A. 2009. Multiple changes in sialic acid biology during human evolution. Glycoconj J. 26: 231–245.

Varki A, Gagneux P. 2009. Human-specific evolution of sialic acid targets: Explaining the malignant malaria mystery? Proc. Natl. Acad. Sci. USA. 106: 14739–14740.

Varki A, Gagneux P. 2012. Multifarious roles of sialic acids in immunity. Ann. N.Y. Aca. Sci. 1253: 16–36.

Varki NM, Varki A. 2007. Diversity in cell surface sialic acid presentations: implications for biology and disease. Lab Invest. 87: 851–857.

Voight BF, Kudaravalli S, Wen X, Pritchard JK. 2007. A Map of Recent Positive Selection in the Human Genome. PLoS. Biol. 4(3): e72.

Wang X et al. 2012. Specific inactivation of two immunomodulatory *SIGLEC* genes during human evolution. Proc. Natl. Acad. Sci. USA. 109: 9935–9940.

Weir BS, Cockerham CC. 1984. Estimating F-Statistics for the Analysis of Population Structure. Evol. 38: 1358–1370.

